# KMT2C knockout generates ASD-like behaviors in mice

**DOI:** 10.1101/2023.05.26.542423

**Authors:** Bastian Brauer, Nicolas Merino-Veliz, Constanza Ahumada, Gloria Arriagada, Fernando J Bustos

## Abstract

Neurodevelopmental disorders have been associated with genetic mutations that affect cellular function, including chromatin regulation and epigenetic modifications. Recent studies in humans have identified mutations in KMT2C, an enzyme responsible for modifying histone tails and depositing H3K4me1 and H3K4me3, as being associated with Kleefstra syndrome 2 and autism spectrum disorder (ASD). However, the precise role of KMT2C mutations in brain disorders remains poorly understood. Here we employed CRISPR/Cas9 gene editing to analyze the effects of KMT2C knockout on animal behavior. Knocking out KMT2C expression in cortical neurons and the mouse brain resulted in decreased KMT2C levels. Importantly, KMT2C knockout animals exhibited repetitive behaviors, social deficits, and intellectual disability resembling ASD. Our findings shed light on the involvement of KMT2C in neurodevelopmental processes and establish a valuable model for elucidating the cellular and molecular mechanisms underlying KMT2C mutations and their relationship to Kleefstra syndrome 2 and ASD.

## INTRODUCTION

Neurodevelopmental disorders are characterized by impairments in brain development that consequently affect behavior, social interactions, communication, cognitive function, and learning abilities, having a significant impact on the life of individuals. Among neurodevelopmental disorders is Kleefstra syndrome 2 that is characterized by intellectual disability, facial dysmorphisms, and autism spectrum disorders (ASD) (Koemans et al., 2017). ASD is a highly variable condition characterized by deficits in social interactions and communication, repetitive behaviors, and restricted interests, which may be associated with comorbid psychiatric, neurological, physical, and/or intellectual disabilities (Lord et al., 2020). Genetic mutations are a prominent factor contributing to the development of ASD. These mutations can lead to the loss of function in a diverse array of genes, resulting in disrupted gene function through mechanisms such as alterations in the reading frame or the creation of premature termination codons (Iossifov et al., 2012; O’Roak et al., 2012; Rubeis et al., 2014; Satterstrom et al., 2020; Tuncay et al., 2022). According to the SFARI GENE database, to date, there are more than 1392 genes associated to ASD that are distributed in all human chromosomes (Banerjee-Basu and Packer, 2010). These genes have many different functions in the cell including neuronal communication, cytoskeleton formation and dynamics, and chromatin regulation and control of gene expression (Iossifov et al., 2012; Rubeis et al., 2014; Stessman et al., 2017; Grove et al., 2019; Ruzzo et al., 2019; Satterstrom et al., 2020). Among chromatin regulating genes, studies using transgenic animals knock-out (KO) for the epigenetic enzymes CHD8, KDM6A or KDM6B show complex behavioral phenotypes that resemble what is observed in humans carrying the mutations in them (Platt et al., 2017; Tang et al., 2017; Gao et al., 2022), establishing model systems to study molecular and cellular mechanisms underlying ASD.

Among the enzymes that control gene expression by chromatin remodeling, and more specifically by altering histone tail modifications is KMT2C also known as MLL3 (Koemans et al., 2017; Eshraghi et al., 2018; Faundes et al., 2018; Satterstrom et al., 2020). KMT2C is a histone lysine methyltransferase enzyme that deposits the H3K4me1 mark, associated with active enhancers, or the H3K4me3 mark related to transcriptionally active regions (Ruthenburg et al., 2007; Jozwik et al., 2016). KMT2C belongs to the COMPASS complex that contributes to essential functions in eukaryotic developmental signaling pathways (Fagan and Dingwall, 2019; Lavery et al., 2020; Cenik and Shilatifard, 2021). Using fruit flies it has been shown that KMT2C binds to promoter regions of genes involved in neuronal processes, and its loss of function produce severe deficits in memory formation (Koemans et al., 2017). In humans, mutations in KMT2C have been found in individuals with intellectual disabilities including Kleefstra syndrome 2 and ASD (Iossifov et al., 2012; Stessman et al., 2017; Lavery et al., 2020; Satterstrom et al., 2020; Dhaliwal et al., 2021; Siano et al., 2022; Tuncay et al., 2022; Zhou et al., 2022). Interestingly only eleven patients have been described with mutations in KMT2C (Siano et al., 2022), with only one long term report showing long lasting phenotypes (Wu and Li, 2022). Thus, phenotypes associated to KMT2C mutants have not been fully characterized.

In this study, we utilized the CRISPR/Cas9 system to generate KMT2C KO models in both cultured cells and mice to investigate the phenotypic consequences of KMT2C loss of function. Using adeno-associated viruses (AAV) as delivery vectors we targeted exon 3 of KMT2C to produce the KO of the gene. As expected, KMT2C KO produced a decrease in the abundance of H3K4me1 and H3K4me3 histone tail marks in cultured neurons. In KMT2C KO mice, we conducted a series of behavioral tests and observed deficits in social interaction, absence of anxiety-like behavior, increased repetitive behaviors, and significant impairments in memory formation.

These phenotypic changes are relevant to the human conditions of ASD and Kleefstra Syndrome 2, both of which are associated with neurodevelopmental disorders that can be attributed to the loss of function of KMT2C. Our findings demonstrate the utility of the CRISPR/Cas9 technology to generate an animal model of KMT2C KO that can be used to investigate the cellular and molecular mechanisms underlying the pathogenesis of KMT2C knockout-mediated disorders.

## Materials and Methods

### Primary neuronal cultures

Postnatal day 0 Cas9 KI mice (C57BL/6J; JAX 026179) were euthanized by decapitation and the whole brain was extracted in ice cold Ca2+/Mg2+-free Hank’s balanced salt solution (HBSS). Meninges were removed, the tissue was minced and incubated with Papain (20 U) for 15 minutes at 37°C. Cells were rinsed twice with HBSS, resuspended by mechanical agitation through fire-polished glass Pasteur pipettes of decreasing diameters, and plated over poly-L-lysine-coated culture plates or cover slips. Cultures were maintained at 37 °C in 5% CO2 in growth media [Neurobasal-A (Life technologies 1088802) supplemented with B27 (Life technologies 17504044), 2 mM L-glutamine (Life technologies 25030-081), 100 U/ml penicillin/streptomycin (Life technologies 15070-063)]. Half of the media was replaced every 3 days. Neuronal cultures were transduced at 3 days in vitro (DIV) using concentrated AAV particles.

### Plasmids

For the expression of CRISPR/Cas9, plasmids were generated in AAV backbones. For sgRNAs 20nt target sequences were selected contiguous to a 5’-NGG photospacer-adjacent motif (PAM). sgRNAs for KMT2C were designed against exon 3 (K1: 5’ GGAAATCAAAGAACAATCTG 3’; K2: 5’ GGAGGATGCTGAAACAGAAG 3’) and cloned into a custom AAV plasmid under the control of the U6 promoter, and coding for tdTomato under the control of the human synapsin1 promoter. For viral packaging we used a plasmid coding for the PHP.eB capsid and pAdDeltaF6 plasmid to express adenovirus E4, E2A and VA genes. pUCmini-iCAP-PHP.eB was a gift from Viviana Gradinaru (Addgene plasmid # 103005 ; http://n2t.net/addgene:103005 ; RRID:Addgene_103005), pAdDeltaF6 was a gift from James M. Wilson (Addgene plasmid # 112867 ; http://n2t.net/addgene:112867 ; RRID:Addgene_112867).

### AAV Production

AAV particles coding for sgRNA against KMT2C under the control of a U6 promoter, together with the red fluorescent protein tdTomato controlled by the hSyn1 promoter were packed using the PHP.eB capsid (Bustos et al., 2017, 2023; Chan et al., 2017). High titer viral particles were purified as described in (Challis et al., 2019). Briefly, HEK 293T were transfected with PEI and PHP.eB capsid plasmids, the vector with KMT2C sgRNA-tdTomato, and the helper plasmid DF6. After 24 h of transfection, the media was replaced for DMEM 1% FBS. 72 h later, medium was collected from the plates and replaced with fresh DMEM 1% FBS. The collected medium was stored at 4ºC. To collect the viruses, 120 h after transfection, the cells were detached from the plate and transferred to 250 mL conical tubes, as well as the collected media. They were centrifuged for 10 min at 2000 g, the supernatant was removed and saved for later use. The pellet was resuspended in SAN digestion buffer (5 mL of 40 mM Tris, 500 mM NaCl and 2 mM MgCl2 pH 8.0) containing 100U/mL of Salt Active Nuclease (SAN) from Arcticzymes and incubated at 37ºC for 1 h. To the supernatant that was saved, a 5x stock solution of 40% PEG 8000 (Sigma) in 2.5M NaCl was added, incubated on ice for 2 h and centrifuged at 4000 g for 30 min in 250 mL bottles. The supernatant was collected and was placed in an Optiprep gradient and ultracentrifuged at 41.000 rpm for 4 h. The phase containing the AAV was rescued and frozen at -80ºC for later use.

### Genomic DNA extraction and T7 endonuclease I assay

Genomic DNA from transduced primary cortical neurons or brain tissue from transduced animals was extracted using Quick-DNA Miniprep Kit (Zymo Research, USA) following manufacturer recommendations. To test KMT2C edition, PCR with primers encompassing the edited region (5’ TACGTTGACCTCAAGGCACAGT 3’, 5’ TAAAACTGTCTCTTGGCCCCCG 3’) were used to determine edition of the locus. PCR products were run in agarose gels and purified by Gel extraction kit (Qiagen). 400ng of gDNA was used for T7 endonuclease assay (NEB Cat#M0302). Assays were run in TBE polyacrylamide gels and visualized using Gel Red.

### RNA extraction and RT-qPCR

RNA was isolated from the tissues and cell cultures using TRIzol (Life Technologies) according to the manufacturer’s instructions as in (Henriquez et al., 2013; Bustos et al., 2017, 2023). To obtain complementary DNA (cDNA), 400 ng of RNA was used. cDNA quantification was performed by qPCR using 3 μL of the cDNA mix, 6 μL Fast Evagreen qPCR Master Mix (Biotium, 31003), 2 μL of nuclease-free water, and 1 μL of 10 mM primers with the program recommended by the maker. The relative abundance was measured by the ddCt method using the GAPDH gene as a control. Transcript detection was performed with specific primers for messenger RNA (mRNA): KMT2C: Fw 5’ TGTTCACAGTGTGGTCAATGTT 3’; Rv 5’ GAGGGTCTAGGCAGTAGGTATG 3’; GAPDH: Fw 5’ ATGGTGAAGGTCGGTGTGAA 3’; Rv 5’ CATTCTCGGCCTTGACTGTG 3’.

### Nuclear protein extraction

Nuclear proteins to determine histone tail modification by western blot were isolated as in (Henriquez et al., 2013). Briefly, cultured cells were washed with cold PBS, centrifuged at 5000 rpm for 5 min and the pellet was resuspended in 5 volumes of cell lysis buffer (50 mM Hepes pH 7.9, 3 mM MgCl2, 20 mM KCl, 0.1% NP-40, 1 mM DTT and Protease Inhibitor Cocktail) and incubated on ice for 10 min. Solution was homogenized with 30 strokes of the pestle and centrifuged at 6000 rpm for 15 min at 4 ºC to separate the cytosolic and nuclear fractions. The pellet was resuspended in 1 volume of Buffer C (10 mM Hepes pH 7.9, 420 mM NaCl, 1.5 mM MgCl2, 25% Glycerol, 0.2 mM EDTA, 1 mM DTT and protease inhibitor cocktail) and incubated for 1 hour. Extracts were sonicated at 50% amplitude in cycles of 30 s ON/ 30 s OFF, centrifuged at 12,000 rpm for 15 min at 4 ºC. Supernatant was recovered and frozen at -80 ºC to be subsequently quantified by Bradford.

### Western blot analysis

Nuclear fractions were separated on polyacrylamide gels and transferred to PVDF membranes (Millipore, USA). Membranes were blocked and incubated overnight at 4C with primary antibodies. After rinsing, the membranes were incubated with secondary antibodies for 30 min at room temperature, rinsed and developed using chemiluminescence (Cell Signaling Technology, USA). Primary antibodies used: H3K4me1 (Diagenode, C15410194), H3K4me3 (Diagenode, C15410003), and H3pan (Diagenode C15410324) as loading control. For detection HRP-conjugated secondary anti-bodies were used (Cell Signaling Technology, USA).

### Animals

All animal procedures and experiments were performed according to the NIH and ARRIVE guidelines and were approved by the animal ethics committee from Universidad Andrés Bello (020/2018). Newborn Cas9 KI mice (C57BL/6J; JAX 026179) were cryoanesthetized in a cold aluminium plate and injected with 1 μL of concentrated AAV (1×10^11^ vg), containing sgRNA or empty vector, in each cerebral ventricle at a depth of 3 mm in the animal’s head at 2/5 of the intersection between lambda and the eye with a 10 μL HAMILTON syringe (Hamilton, 7653-01) and a 32 G needle (Hamilton, 7803-04). After the injection, P0 mice were placed in a heating pad until they recovered their color and temperature, then they were returned to their cage with the mother (Passini and Wolfe, 2001; Kim et al., 2014). 3 weeks after birth, mice from both conditions were weaned off and separated by sex in cages with a 12/12 light/dark cycle with free access to food and water. A chip (p-chips, Pharmseq) was put in the tail of each animal for easy tracking during behavioral test. Behavior tests were performed between 9:00 am and 6:00 pm. At the end of the battery of behavioral tests, the animals were euthanized using isoflurane for subsequent molecular analyses.

### Behavioral overview

All behavioral tests on mice were carried out 8 weeks after AAV injection. Before each test, mice cages were transported to the behavior room and habituated for 30 min in the dark. After completing a test, equipment and devices used were cleaned with 70% ethanol. Tests were recorded and analyzed with ANY-Maze software and/or Graphpad Prism software.

### Rotarod

Motor coordination and capacity was assessed in the Rotarod test. Mice were placed on an elevated accelerating rod for three trials. Each trial lasted for a maximum of 3 min, during which the Rotarod underwent a linear acceleration of 4 rpm per min. Mice weights were registered before the test. Mean time and speed from each animal were registered before falling off.

### Open Field

Mice were tested in an open field (45 × 45 cm) virtually divided into central and peripheral regions with ANY-Maze software. Apparatus were illuminated from above with 300 lux in center and 250 lux in periphery. Animals were allowed to roam freely for 10 min. The total distance traveled and time in center and periphery were analyzed.

### Light and Dark

The apparatus used for this test is a 40 cm box split in half with 390 lux on light side and 0-2 lux on the dark side. Mice were placed in the dark chamber and were allowed to freely explore both chambers for 10 min. Distance traveled, time spent and number of entrances to the light side were analyzed.

### Elevated zero-maze

The apparatus consisted of a 46 cm diameter circular runway and raised 54 cm off the ground. The runway was divided equally into four alternating quadrants of open arcs and closed arcs, with 15 cm walls. Mice started in the center of an open arm and were recorded by video tracking for 10 min. Measures of cumulative open and closed arc times, total open arm entries, distance in each arm and total distance traveled were analyzed.

### Marble burying test

Mice were tested in a 45 × 45 cm box filled with 5 cm deep wood chips and 49 marbles distributed in a 7×7 pattern. Animals were placed in the test cage and allowed to explore and bury the marbles during a 30 min session that was videotaped. At the end of the session the subject was removed and the number of marbles buried (2/3 marble covered by wood chip) was counted.

### Contextual fear conditioning

UGO-BASILE apparatus controlled by ANY-Maze was used. This equipment consisted of a sound attenuating box, fan, light (visible/I.R.), a speaker, a USB camera, a single on-board controller, and a mouse cage. All trials were recorded and all mice underwent a habituation, conditioning and testing phase (Pandian et al., 2020; Bustos et al., 2023). In the habituation (day 1): mice were placed in the fear conditioning cage to explore freely for 5 min and then returned to their cage. During the conditioning phase (day 2): subject mouse was placed in the fear conditioning cage, let explore freely for 2 min and then subjected to an electric shock of 0.75 mA for 2 s. It was allowed to explore freely for 3 min and returned to its cage. On the test phase (day 3): Twenty-four hours after the conditioning phase, the animals were tested for contextual memory. Each mouse was placed in the fear conditioning box, allowed to freely explore for 5 min, and returned to its cage. The number of freezing episodes and freezing time was registered.

### Barnes Maze

A non-reflective gray circular platform (91 cm diameter) with 20 holes (5 cm diameter) evenly distributed along the perimeter, with one hole containing a metal escape tunnel was used. Three exogenous visual cues (length/width ∼30 cm) were used around the platform: black circle, blue triangle and a yellow square. The light was adjusted to 1000 lux in the center of the platform. All animals underwent a phase of habituation, spatial acquisition and testing (Pitts, 2018; Sunyer et al., 2007). For habituation (day 1): each mouse was placed in the center of the platform, directed towards the escape hole, and allowed to remain there for 1 min. Then it was taken and allowed to freely explore the maze for 5 min, and was again allowed to spend 1 min inside the escape hole. If the mouse did not enter within 5 min, it was gently guided near the escape hole selected randomly on the table. During training phase (day 2-4): each animal was introduced into the start box, left in the center of the platform for 10 s and the start box was removed, and simultaneously a 16,000 Hz sound was played. The test ended at 3 min or when the mouse has found the escape hole. This procedure was repeated 2 times per day. During those days the following was recorded: Primary latency: Time to review the escape hole for the first time; Time in the zone of interest; Total distance traveled. In the test phase (day 5): the position of the escape tunnel was changed, and the animal was brought in the start box to the center of the platform, left for 10 s and sound reproduction was started. The test ended at 90 s or when the mouse found the escape tunnel. The number of primary and total errors, primary and total latency, and total distance before finding the gap were recorded. The number of visits to each hole was also measured to show preference.

### Three-Chamber Sociability and Social Novelty Test

This was performed in a transparent acrylic three-chambered apparatus with the following dimensions: 61.5 × 43.5 × 22 cm. Each outer chamber was 20 × 43.5 cm. We used small cages of 8 cm diameter and 18 cm height to put the social (unknown WT mice of the same sex and similar age) and non social stimulus (plastic block of 8 × 4 × 4 cm). A 20 lux illumination was used in this test. On the habituation phase (day 1): stimulus holders were placed in the center of outer chambers. The subject mouse was placed in the central chamber and allowed to explore freely for 10 min. Apparatus and stimulus holders were cleaned between mice with 70% ethanol. During pre-test (day 2): two clean paper balls were prepared and introduced inside each stimulus holder. The mouse was placed in the central chamber and allowed to explore freely for 10 min. Apparatus and stimulus holders were cleaned between mice with 70% ethanol. For the social preference test (day 3): a wild-type mouse was placed in a stimulus holder to be used as a social stimulus and changed for another one every 2 test runs to avoid burnout or social fatigue (Rein et al., 2020). A plastic block was placed in the stimulus holder as a non social stimulus. Once social and non social objects were put in the outer chambers, mouse was placed in the central chamber and allowed to explore freely for 10 min. Behavior was video recorded. Time amount and distance traveled in the social and non social chamber was registered, as well as the interaction time with each stimulus.

### Social Interactions

In this test, an animal from the control condition or KMT2C KO with a wild type animal of the same sex and similar age, were placed in a 30×30×30 cm box for 10 min. Aggressive behavior (e.g. biting, mounting or aggressive grooming) and the amount of social interactions (nose-nose sniffing, nose-face, nose-anogenital area, and grooming) between them were quantified.

### Tube dominance

Tube test apparatus consisted of a smooth transparent acrylic tube of 30 cm length and internal diameter of 2.5 cm. Mice were habituated for 3 days. On day 1, each animal interacted and explored the tube freely for 30 min and was then returned to its cage where a habituation tube was placed with a 10 cm length with an internal diameter of 2.5 cm. A small amount of gel food (Diet Gel Boost) was placed at the end of the cage habituation tube and they were deprived of their common diet food. On day 2, tube inlet was closed and gel food was placed at the end. Animals were allowed to explore freely for 30 min. They were fed and starved again of their common diet for 12h and a small amount of gel food (Diet Gel Boost) was placed at the end of the cage habituation tube. On day 3, the same procedure of Day 2 was repeated, and mice were returned to their common diet. On the training phase (day 4 and 5): mice were taken by the tail and allowed to freely explore for approximately 1 min on the table where the tube was located. Then they were taken from the tail and put on one end of the tube and when the animal entered the tail, it was released. If the animal did not move for several seconds, it was gently prodded with a wooden stick. This step was repeated 5 times per side, so that the mouse passed through the tube a total of 10 times. The same procedure was repeated on day 4 and 5. On the test phase (day 6-9): two mice, one control and one KMT2C KO, were taken and brought to the ends of the tube by the tail. When they entered completely and reached the middle of the tube, their tails were released to begin the confrontation.The test was repeated for 4 days. The mouse that pushes and removes its opponent from the tube was considered the winner and the one that is removed from the tube was considered the loser. All confrontations were video recorded to analyze the times each animal won and lost. The test stopped when the loser had all 4 paws out of the tube. The total number of wins and the number of wins per day were compared between control and KMT2C KO conditions.

### Brain sectioning and mounting

After behaviors to assess brain transduction, animals were deeply anesthetized, and half of the brain was extracted and fixed by immersion PBS + 4% PFA + 4% Sucrose into 30 mL flasks for 24 h. After fixation, a Leica VT1000s vibratome was used to cut 100 μm coronal sections. Slices were kept in PBS and mounted using Fluoromont G (EMS, Hatfield, PA) to preserve the fluorescence signal. Brain images were captured with a Nikon Eclipse TE2000 epifluorescence microscope (Nikon, USA).

### Statistical analysis

Values are presented as mean ± standard error of mean (SEM) for 3 or more independent experiments. Statistical analyzes with Student’s t-test was performed. Values of p<0.05 were considered statistically significant. All statistical analyzes were performed using Graphpad Prism (GraphPad Software Inc.).

## RESULTS

### Gene editing of KMT2C results in knockout in primary culture of cortical neurons and in the mouse brain

To determine the role of KMT2C in the induction of ASD-like behaviors, we used CRISPR/Cas9 technology to knock out gene expression (Cong et al., 2013; Swiech et al., 2014). First, we designed sgRNAs targeting the coding sequence of KMT2C. We targeted exon 3 of the gene since it is a common exon for all splice variants (Figure 1A). Cortical neuron cultures of Cas9 KI mice were transduced using AAVs coding for TdTomato alone as a control or together with sgRNA K1 or K2. Ten days after transduction, genomic DNA was extracted, and the T7 endonuclease I assay was performed. We observed that both sgRNAs targeting KMT2C exon 3 were able to edit the genomic locus, shown by the smaller size DNA bands (Figure 1B). Additionally, total RNA was extracted, and RT-qPCR was performed to determine changes in KMT2C expression. We observed that both K1 and K2 sgRNAs significantly reduced KMT2C expression by more than 60% in primary cortical neurons (Figure 1C). KMT2C is a component of the COMPASS chromatin remodeling complex that can increase the histone tail modifications H3K4me1 and H3K4me3 at enhancers and promoters to induce gene expression (Jozwik et al., 2016). Therefore, we tested the presence of these histone tail marks in nuclear extracts of transduced cortical neurons. We found that the transduction of neurons with K1 or K2 decreased the total abundance of H3K4me1 and H3K4me3 (Figure 1D). This data was quantified showing a significant reduction in the relative expression over H3 of both H3K4me3 (Figure 1E) and H3K4me1 (Figure 1F). This data shows that the CRISPR/Cas9 system can edit the KMT2C genomic locus, reduce its expression, and decrease the presence of histone tail modifications associated with its function in KMT2C KO cultured neurons.

**Figure 1.**
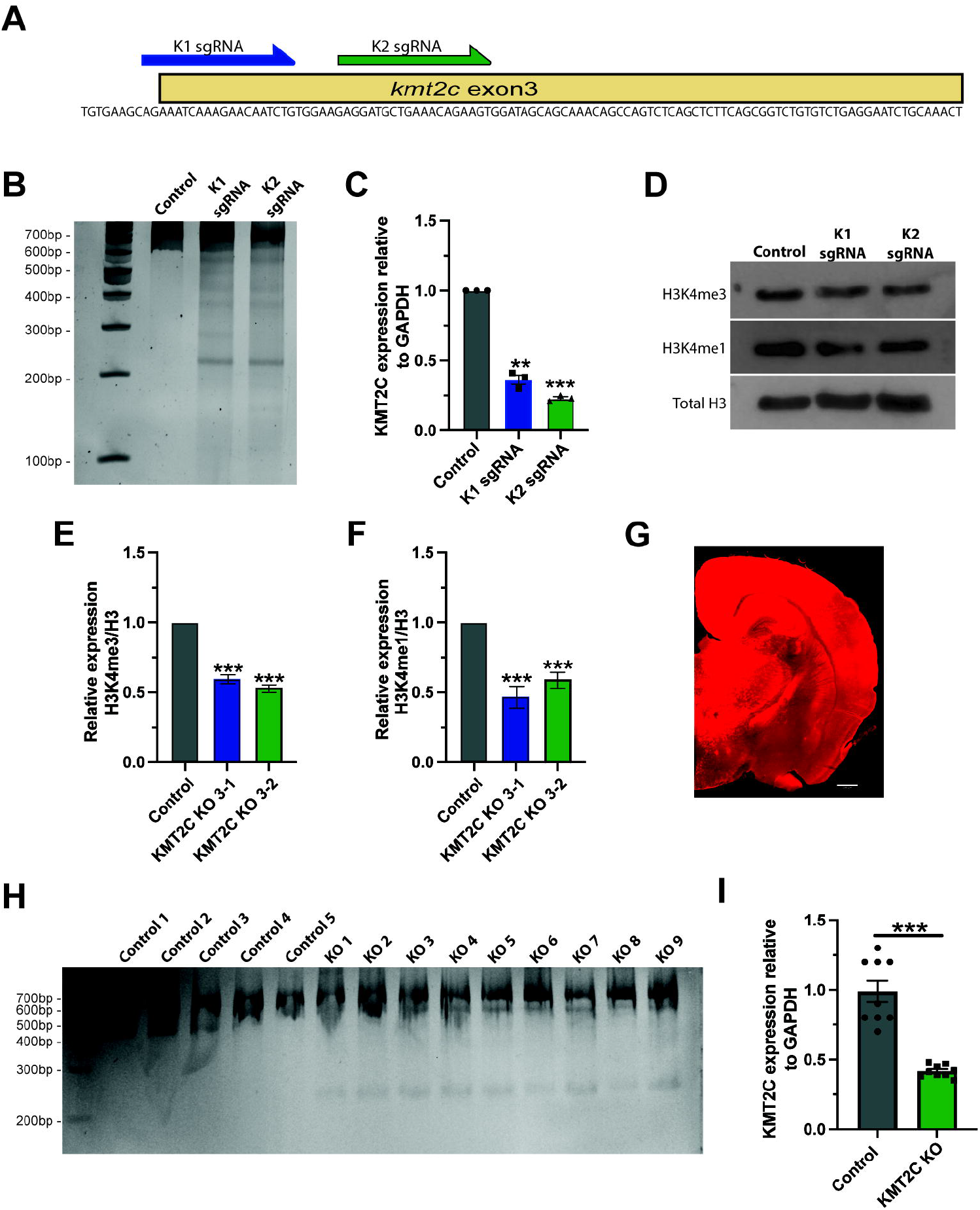
Knockout of KMT2C by gene editing *in vitro* and *in vivo*. **(A)** Scheme showing the exon 3 of KMT2C genomic sequence and the positions where K1-K2 sgRNA were designed. **(B)** T7 endonuclease I assay from transduced cultured neurons. **(C)** RT-qPCR to determine expression levels of KMT2C relative to GAPDH in transduced cultured neurons. **(D)** Representative image of western blot analysis of total H3 histone, H3K4me3, HeK4me1. (E-F) Quantification of the relative expression of H3K4me3 **(E)** or H3K4me1 **(F). (G)** Representative image of brain slice after transduction by intracerebroventricular injection of CRISPR/Cas9. **(H)** T7 endonuclease I assay from transduced cortical tissue of injected animals. **(I)** RT-qPCR to determine expression levels of KMT2C relative to GAPDH in transduced cortical tissue of injected animals. Bars represents mean ±SEM; **p<0.01, ***p<0.001. Students t-test was used to determine significance compared to wild-type condition. Scale bar = 500µm.

In humans, loss-of-function mutations in KMT2C have been associated to the appearance of ASD-like behaviors and Kleefstra syndrome 2 (Koemans et al., 2017; Satterstrom et al., 2020; Siano et al., 2022; Wu and Li, 2022), however limited information is available about the phenotype. To determine whether KO of KMT2C in mice can produce ASD-like behaviors, we used AAVs to express the previously characterized K2 sgRNA in Cas9 knock-in animals. We selected K2 sgRNA from our culture experiments because it showed a more significant reduction in mRNA expression compared to control neurons and K1 (Figure 1C). The fluorescence reporter TdTomato was used to show transduction efficiency. AAVs were packed using the PHP.eB capsid (Chan et al., 2017) to efficiently transduce the whole brain after injection into the cerebral ventricles of P0-P1 animals (Bustos et al., 2023). At eight weeks, animals were subjected to behavioral testing. To be included in the final behavioral analyses, animals needed to meet three parameters after euthanizing and dissecting their brains: 1) a strong fluorescence signal widely spread in the brain (Figure G); 2) the T7 endonuclease I test showed gene editing (Figure 1H); and 3) RT-qPCR analyses showed a >50% reduction in KMT2C expression levels (Figure 1I). A total of 9 animals met these criteria, and the following results for KMT2C CRISPR injected animals (henceforth KMT2C KO mice) are only based on these 9 animals and 15 animals used as control. Fluorescence imaging of brain sections, show high and broad expression of tdTomato (Figure G). All KMT2C KO (KO1-9) animals show lower size bands in the T7 endonuclease I assay, showing that gene editing was successful (Figure 1H, KO1-9) compared to 5 representative control animals (Figure 1H, Control 1-5). In addition, RT-qPCR from brain tissue showed >50% reduction in the expression levels of KMT2C in injected animals (Figure 1I). These results demonstrate that CRISPR/Cas9 system can edit the KMT2C genomic locus and reduce its expression in the mouse brain.

### KMT2C KO animals exhibit no signs of behaviors associated with anxiety

Eight weeks after injection, KMT2C KO and wild-type littermate (Control) animals underwent behavioral testing. Importantly, KMT2C knockout did not result in any discernible differences in growth. To assess whether KMT2C KO animals had any locomotion deficits that could impact their performance in the upcoming battery of behavioral tests, the rotarod test was performed. KMT2C KO animals did not exhibit any locomotion difficulties compared to their wild-type littermates, as both groups spent similar amounts of time on the rotarod apparatus (Figure 2A). One of the hallmarks of both human and animal ASD cases is the manifestation of anxiety-like behaviors (Silverman et al., 2010; Lord et al., 2020; Pandian et al., 2020; Bustos et al., 2023). To assess whether knockout of KMT2C could produce similar behaviors in mice, we subjected KMT2C KO animals to behavioral analyses after 8 weeks. In the open field test, animals were allowed to explore for 10 min, and the time spent in the center or periphery of the apparatus and the distance traveled was measured (Figure 2B). We observed no significant differences between KMT2C knockout animals and control littermates in total distance traveled (Figure 2C), time spent in the center (Figure 2D), or time spent in the periphery (Figure 2E). However, we when we looked in more detail at the time spent in the center, we found significant reduction in the distance travelled in the center by KMT2C KO animals (Figure 2F), suggesting appearance of anxiety since animals freeze in the zone.

**Figure 2.**
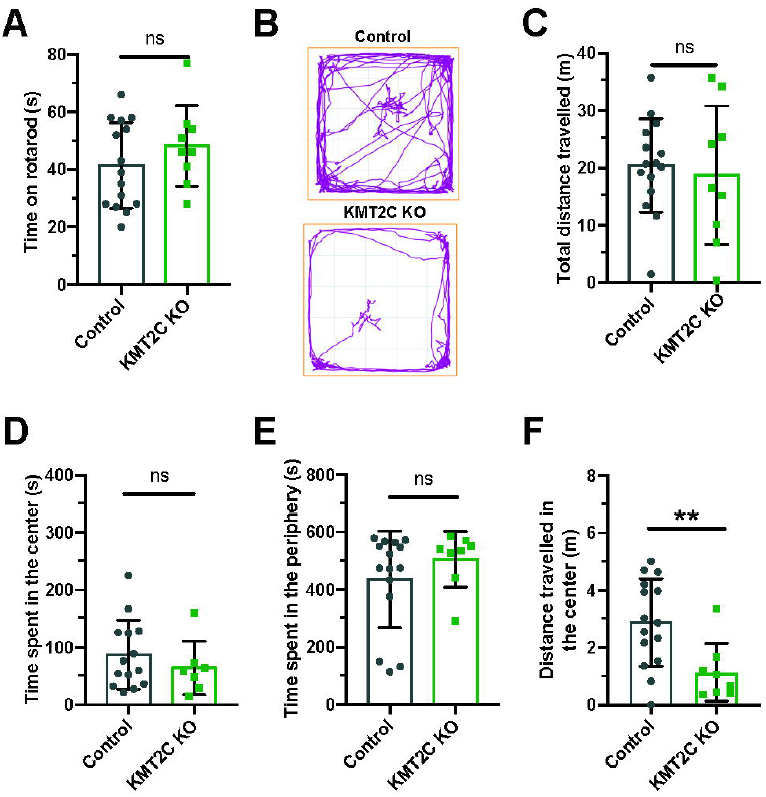
No signs of anxiety behaviors are observed in KMT2C KO animals. **(A)** Time on the rotarod apparatus. **(B)** Representative trace plots of open field test in Control and KMT2C KO animals. **(C-F)** Quantification of the total distance travelled **(C)**, time spent in the center area **(D)**, time spent in the periphery **(E)**, and the distance travelled in the center of the open field arena **(F)** Bars represents mean ±SEM; **p<0.01. Students t-test, n=14 Control animals and n=9 KMT2C KO animals.

Then animals were tested in the light-dark apparatus, where they were placed in the dark chamber and allowed to explore for ten minutes. We observed no significant difference in the number of crosses to the light compartment (Figure 3A) or time spent in the illuminated field (Figure 3B). Similarly, in the elevated zero maze, KMT2C KO animals did not differ significantly from wild-type littermates in the time spent in the open zone (Figure 3C) or distance traveled (Figure 3D) after exploring the platform for ten minutes. Finally, to determine whether KMT2C KO animals showed signs of anxiety or repetitive behavior, we used the marble burying test, in which KMT2C KO and Control animals were placed in a box with 49 marbles to be buried in a thirty-minute interval. We found that KMT2C KO animals buried more marbles than Control littermates within the given timeframe (Figure 3E). Taken together, our data show that KMT2C KO animals exhibit repetitive behaviors, as observed in the marble burying test, and display almost no signs of anxiety-like behaviors that could be measured using these behavioral tests.

**Figure 3.**
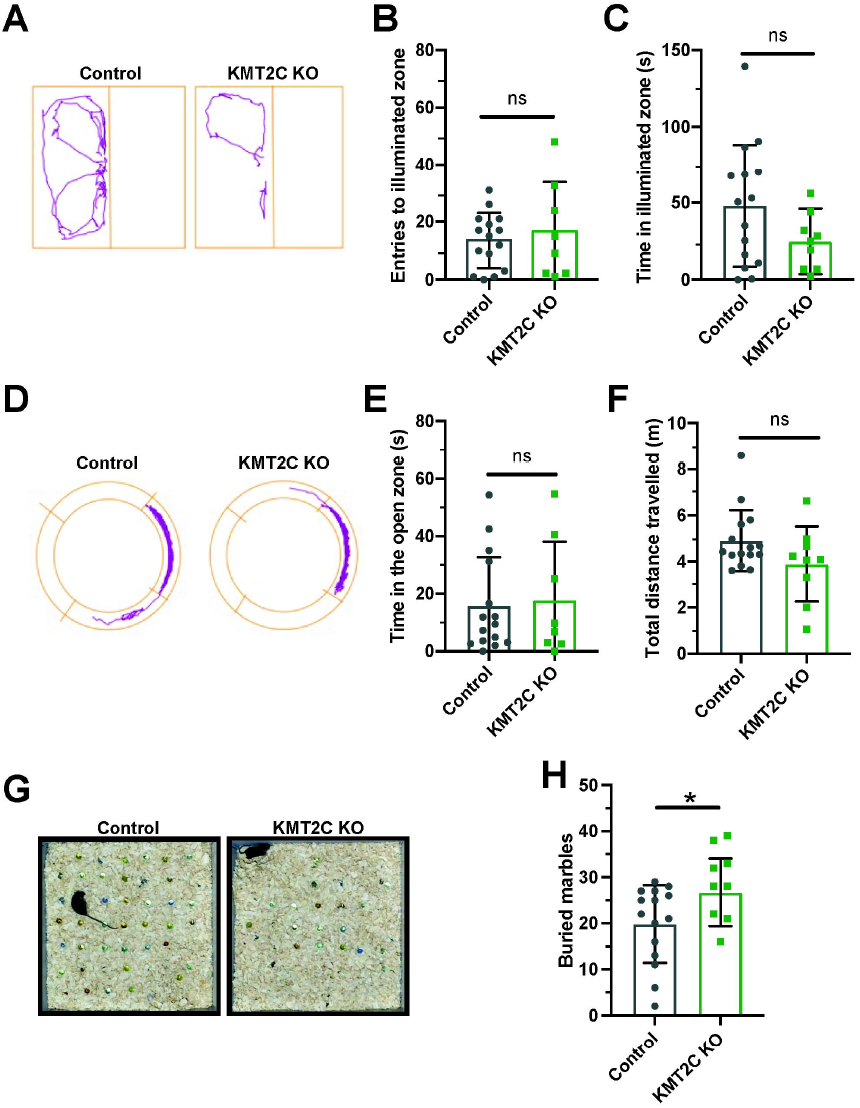
KMT2C KO animals evidence repetitive behaviors. **(A)** Representative trace plots of dark and light test. **(B-C)** Quantification of the number of entries to light zone **(B)**, and the time spent in the light in the dark and light apparatus. **(D)** Representative trace plots of the zero-maze test. **(E-F)** Quantification of the time spent in the open zone **(E)**, and the total distance travelled **(F)** in the zero maze. **(G)** Representative images of the final number of marbles buried by Control and KMT2C KO animals. **(H)** Quantification of the number of marbles buried during the marble burying test. Bars represents mean ±SEM; *p<0.05. Students t-test, n=14 Control animals and n=9 KMT2C KO animals.

### KMT2C KO animals show impaired social behaviors

Individuals with ASD often exhibit difficulties in social interaction, which can manifest as a lack of aptitude or skill in performing social interaction (Silverman et al., 2010; Lord et al., 2020). To assess the ability of KMT2C KO mice to engage in social interaction, we employed the three-chamber social interaction test leaving the animal to explore for ten minutes (V. et al., 2007; B. et al., 2010; Pandian et al., 2020; Bustos et al., 2023). KMT2C KO and Control animals spent similar time in the novel mouse region (NM) (Figure 4A) and novel object region (NO) (Figure 4B). However, when the time spent directly interacting with NM or NO was analyzed in more detail, we found that KMT2C KO animals spent significantly more time engaging directly with the NM (Figure 4C) and NO (Figure 4D) than their wild-type littermates. This suggests that KMT2C KO mice exhibit alterations in social behavior compared to Control mice.

**Figure 4.**
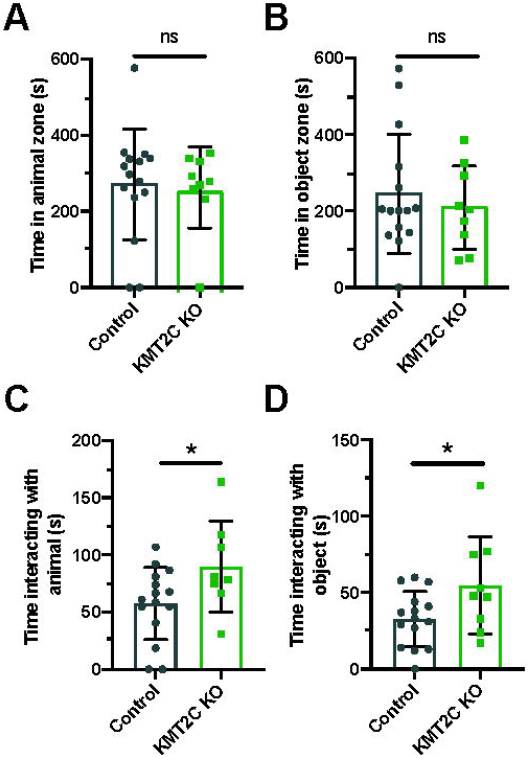
Impaired social behaviors of KMT2C KO animals in the 3-chamber test. **(A-D)** Quantification of the time spent in the animal’s zone **(A)**, the time spent in the object zone **(B)**, the time spent interacting directly with the animal **(C)**, and the time interacting directly with the object **(D)** Bars represents mean ±SEM; *p<0.05. Students t-test, n=14 Control animals and n=9 KMT2C KO animals.

To gain insight into the social behavior observed, we used the social interaction test. Pairs of animals of the same sex from KMT2C KO and Control animals were let to interact in a clear box for 10 minutes and recorded to quantify interactions. The quantification of social sniffing in animal pairs showed no significant difference in the frequency of nose-nose (N-N) interactions (Figure 5A) or nose-head (N-H) interactions (Figure 5B). However, we observed a significant decrease in the number of nose-anogenital (N-A) interactions in KMT2C KO mice (Figure 5C), indicating a decrease in social investigation activity. Additionally, we found that KMT2C KO animals displayed a significant increase in self-grooming episodes compared to their wild-type littermates (Figure 5D). Lastly, we used the tube dominance test to determine social hierarchy and interaction among animals. Results show that KMT2C KO animals won a significantly greater number of times against wild-type animals in the test tube (Figure 5E). This data demonstrates that KMT2C KO animals exhibit social impairments, as evidenced by a decrease in social investigation, an increase in self-grooming episodes to avoid interactions, and avoidance of direct contact with other animals. These behaviors suggest that KMT2C KO animals have difficulty engaging in social interactions and may display aggression when faced with such situations.

**Figure 5.**
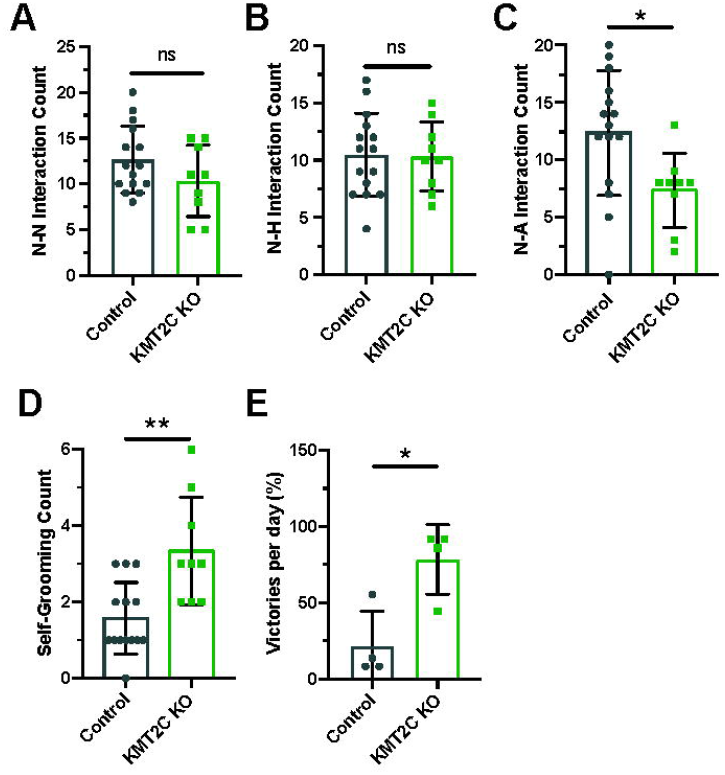
KMT2C KO animals show impaired social interaction beahviors. **(A-C)** Quantification of the **(A)** nose-nose (N-N), **(B)** nose-head (N-H), and **(C)** nose-anogenital (N-A) interactions between Control or KMT2C KO animals. **(D)** Quantification of the self-grooming behavior shown by Control or KMT2C KO animals. **(E)** Quantification of the number of victories in the tube dominance test. Bars represents mean ±SEM; *p<0.05, **p<0.01. Students t-test, n=14 Control animals and n=9 KMT2C KO animals.

### KMT2C KO mice exhibit severe memory formation deficits

Cognitive impairments are a crucial feature of ASD phenotypes. Human with mutations in KMT2C have shown to have mild to severe intellectual disability, phenotype also associated to ASD (Koemans et al., 2017; Siano et al., 2022; Wu and Li, 2022). To determine whether KMT2C KO animals have impaired capacity to form memories, we utilized two behavioral tests. First, we employed the contextual fear conditioning paradigm to analyze long-term memory formation after 24 hours. Animals were tested for five minutes, and we observed that KMT2C KO animals froze for a significantly reduced amount of time compared to Control animals (Figure 6A). Next, we used the Barnes maze apparatus to assess spatial learning and memory (Pitts, 2018; Gawel et al., 2019). Similarly, to what is observed in the open field apparatus, no significant difference in the total distance travelled between KMT2C KO and controls animals (Figure 6B). However, when we quantified the primary latency - time to reach the escape hole for the first time - we observed a significant increase in the time required by KMT2C KO animals, and some of them never reached the escape hole (Figure 6C). Finally, to assess whether KMT2C KO animals showed deficits in spatial memory, we moved the escape hole to a new location and quantified the time spent in the region where the original escape hole had been located. We observed a significant decrease in the time spent in the target region by KMT2C KO animals compared to Controls (Figure 6D). Taken together, our data suggests that KMT2C KO animals exhibit reduced memory formation capacity and significant cognitive impairments in both behavioral tests used.

**Figure 6.**
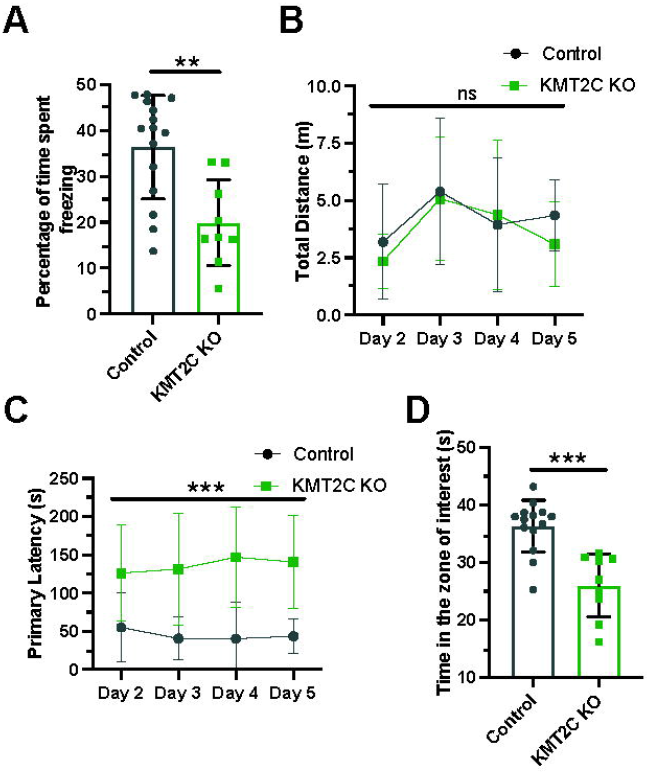
Memory formation is significantly impaired in KMT2C KO animals. **(A)** Percentage of time spent freezing by Control and KMT2C animals in the contextual fear conditioning test. **(B)** Quantification of the total distance travelled in the Barnes maze apparatus. **(C)** Primary latency of Control and KMT2C KO animals in the Barnes maze apparatus. **(D)** Quantification of the time spent in the zone of interest where the escape hole was formerly located. Bars represents mean ±SEM; **p<0.01, ***p<0.001. Students t-test, n=14 Control animals and n=9 KMT2C KO animals.

## DISCUSSION

Neurodevelopmental disorders, including ASD and Kleefstra Syndrome 2, have been associated with mutations in genes involved in chromatin regulation (Rubeis et al., 2014; Eshraghi et al., 2018; Satterstrom et al., 2020; Chen et al., 2021; Siano et al., 2022). Some of these genes encode enzymes responsible for modulating histone tail modifications, which play a crucial role in gene expression regulation (Bannister and Kouzarides, 2011).

In this study, we utilized CRISPR/Cas9 technology to disrupt the expression of KMT2C, an H3K4 methyltransferase enzyme, both *in vitro* and *in vivo*. Our results demonstrate a significant reduction in KMT2C expression in vitro, accompanied by a concomitant decrease in the overall levels of H3K4me1 and H3K4me3, histone tail modifications associated with the function of KMT2C (Shilatifard, 2012; Faundes et al., 2018). This reduction in histone tail marks indicates the successful gene editing of KMT2C. Furthermore, the changes in these histone tail marks suggest a mechanism by which KMT2C KO may impact gene networks involved in the observed phenotype. To further elucidate these alterations, future ChIP-Seq experiments will be conducted to examine changes in histone marks associated with promoters and analyze the specific genes and gene networks involved in the manifestation of the phenotype.

In humans, studies have demonstrated a strong association between mutations in KMT2C and carcinogenesis, highlighting the connection between epigenetic regulation and the development of cancer (Fagan and Dingwall, 2019). However, the understanding of the impact of KMT2C mutations on brain disorders remains limited. In humans, loss of function mutations of KMT2C causes ASD and Kleefstra syndrome 2, characterized by intellectual disability and ASD-like behaviors (Koemans et al., 2017). Only eleven cases of Kleefstra syndrome 2 have been reported in literature, thus not much evidence on the behavioral phenotype is described (Koemans et al., 2017; Chen et al., 2021; Siano et al., 2022; Wu and Li, 2022). Using AAV intracerebral ventricular injections at postnatal day 1, we injected CRISPR/Cas9 to produce KMT2C KO. We observed a robust decrease in the expression of KMT2C following gene editing *in vivo*, which enabled us to investigate the behavioral changes associated with ASD. We subjected the animals to a battery of test that are design to test ASD like behaviors (Pandian et al., 2020; Bustos et al., 2023). Similar experiments have been conducted to characterize the behavioral phenotypes in mutant animals for KMD6A and KMD6B (Tang et al., 2017; Gao et al., 2022). KMT2C KO animals did not show signs of affected growth or locomotor abilities, thus effects on the posterior tests are not due to locomotive problems. Using the open field test, light and dark apparatus, and elevated zero maze, we did not observe significant differences in anxiety-related behaviors. This observation is supported by the phenotypes observed in humans carrying KMT2C mutations where anxiety is not described as a main characteristic of the individuals (Koemans et al., 2017; Grove et al., 2019; Siano et al., 2022; Wu and Li, 2022). Repetitive behaviors are a recognized characteristic of ASD phenotypes, as evidenced by studies conducted on other mouse models (Silverman et al., 2010; Lord et al., 2020; Pandian et al., 2020; Bustos et al., 2023). To assess repetitive behaviors in KMT2C KO animals, we employed the marble burying test and quantified grooming behaviors. Our results indicate that KMT2C animals exhibit increased repetitive behaviors. While this specific characteristic has not been described in humans with KMT2C mutations, it is important to note that repetitive behaviors are complex and may be associated with the presence of comorbid conditions such as attention-deficit/hyperactivity disorder (ADHD), as observed in individuals with KMT2C mutations (Wu and Li, 2022).

In the social tests performed, KMT2C KO animals displayed increased social interactions and demonstrated dominance over WT animals in the tube dominance test. While this specific phenotype has not been described in humans due to limited available data, it highlights a novel observation that warrants further investigation in individuals with KMT2C mutations. By exploring this phenotype in human subjects with KMT2C mutations, we may gain valuable insights into the role of KMT2C in social behavior and its potential implications for neurodevelopmental disorders. Intellectual disability is a well-documented phenotype observed in all human patients with KMT2C mutations, as supported by previous studies (Koemans et al., 2017; Siano et al., 2022; Wu and Li, 2022). To determine whether KMT2C KO animals exhibit impairments in memory formation, which is closely associated with intellectual disability, we conducted the fear conditioning and Barnes maze tests. Results revealed severe deficits in memory formation in KMT2C KO animals, with some individuals failing to learn the required tasks altogether. This finding aligns with the phenotypes observed in humans with KMT2C mutations and suggests that KMT2C plays a critical role in cognitive processes, including memory formation, thereby contributing to intellectual disability.

To the best of our knowledge, this study represents the first comprehensive characterization of the behavioral phenotype of KMT2C KO animals. Particularly, using CRISPR/Cas9 gene editing we were able to KO KMT2C in the entire brain to assess the behavioral phenotype. The phenotypic similarities observed between KMT2C KO animals and humans with KMT2C mutations indicate the relevance and translational potential of our approach. This animal model wil allow to investigate the underlying cellular and molecular mechanisms involved in the development of ASD and Kleefstra syndrome 2 associated with KMT2C mutations.

In summary, our study contributes to the understanding of KMT2C-related neurodevelopmental disorders. In addition, the generated animal model offers a promising avenue for investigating therapeutic interventions and advancing our knowledge of the molecular pathways involved in ASD and Kleefstra syndrome 2.

## ACKNOWLEDGEMENTS

This work was supported by: ANID Fondecyt Iniciacion 11180540 (FJB), ANID PAI 77180077 (FJB), ANID-FONDECYT 1220480 to GA, UNAB DI-02-22/REG (FJB), DI-03-21/APP UNAB (BB). We thank Lorena Varela-Nallar for critical reading of the manuscript and helpful feedback.

